# DeepGraphMut: A graph-based deep learning method for cancer prognosis using somatic mutation profile

**DOI:** 10.1101/2024.12.03.626568

**Authors:** Aswin Jose, Akansha Srivastava, P.K. Vinod

**Affiliations:** Centre for Computational Natural Sciences and Bioinformatics, IIIT, Hyderabad-500032, India

## Abstract

Cancer remains a leading cause of morbidity and mortality worldwide. Despite advances in genomics, identifying clinically relevant subtypes of cancer remains challenging due to its complex and heterogeneous nature. In this work, we propose DeepGraphMut (DGM), a novel graph-based deep-learning pipeline that integrates somatic mutation data with protein-protein interaction (PPI) networks. By employing a graph autoencoder with a graph attention layer and a node-level attention decoder, DGM generates patient-specific clinically relevant encodings for unsupervised and supervised tasks. We demonstrate the effectiveness of DGM across 16 cancer types comprising of 7352 samples from The Cancer Genome Atlas (TCGA). Unsupervised clustering reveals distinct subtypes with significant survival differences in 11 cancer types. In supervised analysis using a Cox regression model, DGM demonstrates excellent performance in predicting survival outcomes, achieving a high concordance index (c-index) value in the range of 0.7 across most cancers, underscoring its robust predictive performance using only somatic mutation data. Furthermore, DGM outperforms its lightweight variant and the network-based stratification method in both unsupervised and supervised analyses. In summary, this study presents a promising approach for cancer subtype identification and prognosis, especially in resource-limited settings where multi-omics data may not be readily available. By leveraging the strengths of graph learning and network biology, DGM offers a valuable tool for advancing personalized medicine.

## Introduction

According to GLOBOCAN, there were close to 20 million new cancer cases and 9.7 million cancer deaths worldwide in 2022, with approximately one in five people developing cancer^1^. Traditionally, cancer classification relies on the site of origin and histological characteristics^2^. The current treatment landscape is often constrained by a one-size-fits-all approach^3,4^ due to an incomplete understanding of the underlying molecular and regulatory changes driving cancer progression. While cost-effective and the best option available given current knowledge, this strategy is far from ideal for addressing the diverse and complex nature of cancer. It is important to use molecular biomarkers for cancer stratification to help guide personalized treatment strategies and improve patient outcomes.

Cancer cells evade signals that control cell behaviour due to various DNA abnormalities, such as somatic mutations, alterations in copy number, and changes in DNA methylation patterns^5^. The advent of high-throughput sequencing (HTS) technology and its increasing accessibility have revolutionized our understanding of the human genome, ushering in a new era of data abundance^6^. Somatic mutation profiles, obtained by comparing the genome or exome of a patient’s tumour to that of the germ line using high-throughput sequencing, are a promising source of data for cancer stratification. These profiles are presumed to contain the causal drivers of cancer progression^7^. However, somatic mutation profiles are extremely sparse and heterogeneous, making stratification challenging.

Network Biology enables the interpretation and modelling of complex biological systems through the integration of omics data and biological interactions^8,9^. Recently, the application of biological networks has proven instrumental in unravelling biological mechanisms, understanding disease origins, and forecasting responses to therapies at both molecular and systemic levels^10^. Network-based approaches, particularly network diffusion(ND) or propagation, have gained prominence in analyzing HTS datasets by leveraging known or inferred gene relationships. The network-based stratification (NBS) method proposed by Hofree et al. (2013) is widely utilized to integrate protein-protein interaction (PPI) network with tumor mutation data^11^. Patient encodings are obtained by propagating mutational information across the PPI network. Based on these encodings, patients are further stratified into clinically relevant groups^11^. Zong et al. (2015)^12^ applied NBS to stratify 13 cancers using gene panels, discovering survival-related subtypes in five of them, while PyNBS further refined the NBS approach by utilizing a more compact PPI network for improved outcomes^13^. Other works have integrated different genomic data profiles with PPI networks at various stages, classified based on the point of data integration (ND first, ND during, and ND after)^14^. Additionally, the Network embedding method has been introduced as another approach for patient stratification. Notably, the Network Embedding Stratification (NES) approach combines the human PPI network with genome-scale somatic mutation profiles, hypothesizing that patients with mutations in similar network regions are more likely to be of the same subtype. This method involves gene vectorization through network embedding using struc2vec, patient feature construction by integrating somatic mutation profiles with gene vectors, and patient stratification using machine learning approaches^15^.

A step further from network diffusion, graph learning involves applying machine learning techniques to graph-structured data by transforming graph characteristics into feature vectors of uniform dimensions within an embedding space. This allows for direct processing of graph data by the model without the need for dimensionality reduction. These methods, predominantly rooted in extending deep learning technologies, excel at encoding graph structures into vectorial representations, with the resulting vectors occupying a continuous vector space and aiming to extract relevant graph features for further analysis^16^. These techniques substantially enhance the precision and depth of insights that can be obtained from graph data. They showcase the advanced capabilities of graph learning in refining our understanding and interpretation of complex network structures, surpassing the limitations of traditional network diffusion approaches. In recent years, various approaches have been developed to understand omics data through graph-based deep learning^17^. A notable stratification approach is the Consensus-guided Graph Autoencoder (CGGA) method, which effectively identifies cancer subtypes by integrating structure information and node features through graph autoencoders and iteratively refining feature learning with omic-specific similarity matrices^18^. Also, the approach omicsGAT, which leveraged the self-attention mechanism, generated embeddings from gene expression data by connecting samples with similar features, indicating potential similarities in disease outcomes or cell types. It incorporates network information through an adjacency matrix and employs a multi-head framework to assign varying attention levels to each neighbour of a sample, capturing their differential contributions to the sample’s embedding^19^. These existing graph-based deep learning approaches primarily focus on node-level tasks, beginning with the creation of a patient similarity network and then applying either supervised or unsupervised analysis.

In this work, we introduce a novel graph-level framework, DeepGraphMut(DGM), leveraging a graph autoencoder equipped with a graph attention layer and a decoder featuring node-level attention to generate clinically relevant patient-wise encodings in an unsupervised manner. This approach integrates prior biological knowledge, such as PPI networks, with somatic mutation information for cancer subtype identification and prognosis, harnessing the power of graph learning, modern computational capabilities, and data derived from HTS technologies. By treating each patient as an individual graph, our method enables a comprehensive and personalized analysis of the data.

## Results

### Overview of the pipeline

Figure 1 illustrates the comprehensive workflow of the DGM pipeline. This graph-based deep learning pipeline integrates somatic mutation data and PPI network to learn meaningful representations of genes for identifying cancer subtypes and predicting patient survival. The process begins by preparing the input data, which involves deriving a cancer-specific subnetwork from the human PPI network (see Methods). This subnetwork is constructed from PCNet, using genes sourced from the Network of Cancer Genes (NCG)^20^. A patient-specific network is then created by combining this NCG-derived subnetwork with somatic mutation data, representing each patient as an individual network with mutation profiles encoded as node values. Edge connections are kept consistent across patients, allowing for a stable network structure while capturing each patient’s unique mutational landscape. This network is then processed through the DeepGraphMut (DGM) pipeline, which includes encoding and decoding modules within an autoencoder framework. In the encoding module, GraphNorm is applied to optimize the normalization procedure by leveraging graph structure information. This technique offers notable advantages over traditional normalization methods, particularly in managing heterogeneous data types and complex network structures. In the decoding module, rather than using a traditional edge decoder, we employ a node decoder that focuses on learning variations in node values, representing patient-specific mutation profiles. This design captures critical differences in mutation impacts across patients. The node decoder transforms the latent representation back into a series of node values, which are directly compared with the original omics data input to the encoder. This node-focused approach enables the extraction of salient features from the latent space, enhancing the analysis of the omics data. The model is trained until the optimal weights are learned, and the weights corresponding to the minimal validation loss are used for generating the encodings of input data. The encoded data then undergoes processing through a mean pooling layer, facilitating subsequent analyses using both supervised and unsupervised approaches. The DGM pipeline was used to analyze somatic mutation data from 16 different cancer types in the TCGA (see Supplementary information).

**Figure 1.**
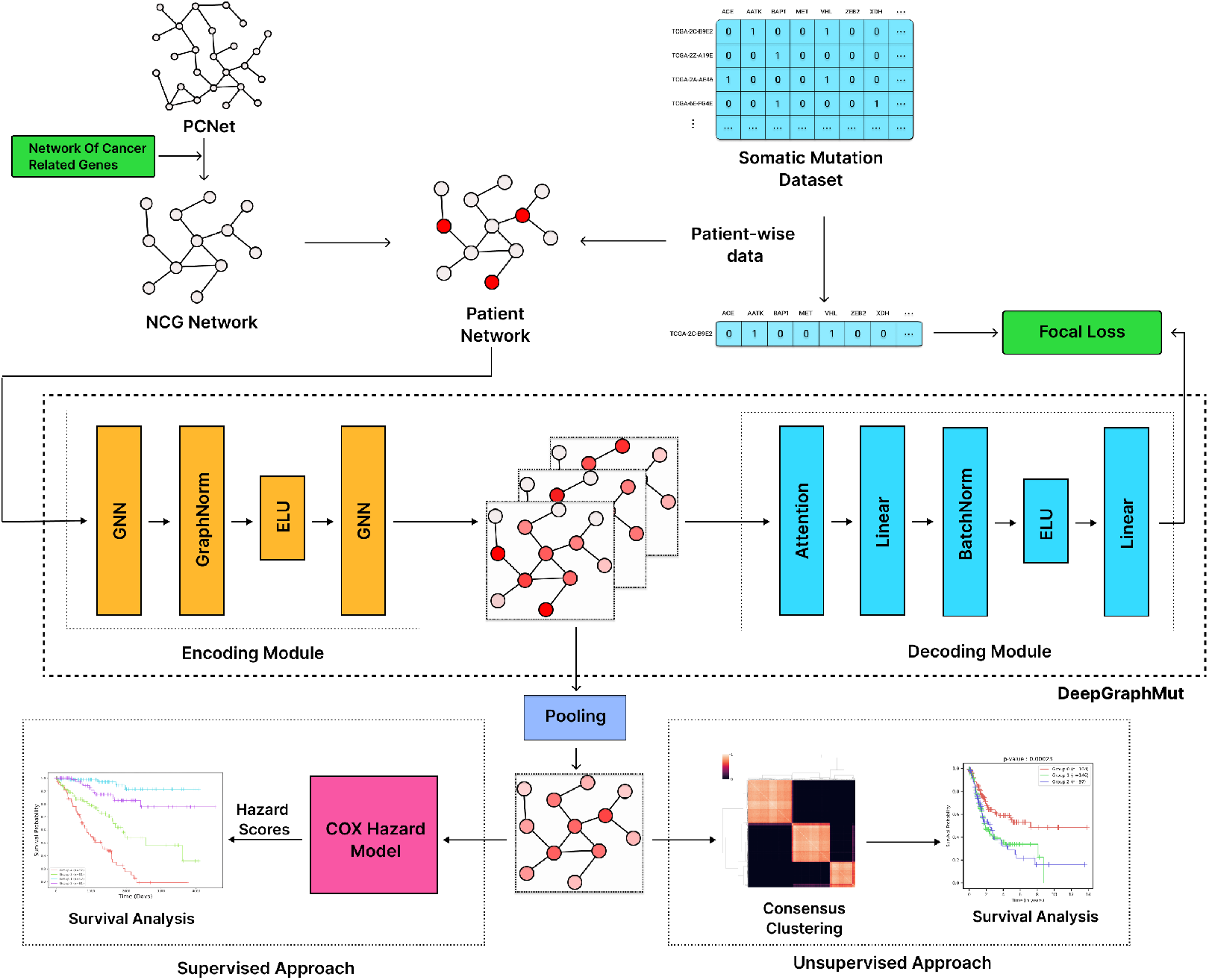
Overview of DeepGraphMut pipeline

### Stratification of patients into subtypes with distinct survival outcomes

The gene representations learned from the DGM were used for clustering patients. Consensus clustering was applied to these representations to stratify cancer patients into distinct subtypes. We employed two clustering algorithms: Partitioning Around Medoids (PAM) and K-means. Survival analysis was then performed to evaluate the differences in survival among the identified cancer subtypes. The optimal number of clusters for each cancer type was determined based on silhouette scores, cophenetic correlation coefficients, and p-values. The DGM pipeline consistently performed well in stratifying patients into subtypes across the 11 cancers studied. Each subtype included a substantial number of patients, exhibited high cophenetic correlation coefficients, and yielded significant p-values (Table 1 and Figure 2). These results highlight the pipeline’s broad applicability and effectiveness in addressing diverse cancer types. The PAM clustering algorithm demonstrated superior stability and statistical significance in clustering across most datasets. Notably, PAM achieved a balanced distribution of patients within each cluster, enhancing the interpretability and potential clinical relevance of the clustering outcomes (Table S1). In contrast, while K-means clustering identified some significant clusters, it often resulted in less distinct or significant groupings compared to PAM. An exception was noted in the analysis of GBM, where K-means outperformed PAM by producing significant clusters. Additionally, the optimal number of clusters varied across the 11 cancer types: two subtypes were identified for KIRC, GBM, and UCEC; three subtypes for BLCA, HNSC, OV, and STAD; four subtypes for LIHC; and five subtypes for LGG, LUSC, and SKCM (Table 1). The Kaplan-Meier plots illustrate the differences in survival probabilities among the cancer subtypes (Figure 2). Notably, for BLCA, GBM, LGG, and SKCM, the observed p-values were very low, indicating highly significant differences in survival outcomes of the identified subtypes. The consensus maps and the corresponding Kaplan-Meier plots for the other cancers are provided in the Supplementary Material (Figure S1).

**Table 1.** Unsupervised clustering results on encodings obtained using the DGM model with NCG network.

| Cancer | Clustering Algorithm | Clusters | Silhouette Score | p-value | CCC |
| --- | --- | --- | --- | --- | --- |
| BLCA | pam | 3 | 0.45 | 0.00023 | 0.99 |
| GBM | kmeans | 2 | 0.06 | 0.00013 | 0.995 |
| HNSC | pam | 3 | 0.42 | 0.02546 | 0.985 |
| KIRC | pam | 2 | 0.35 | 0.0168 | 0.996 |
| LGG | pam | 5 | 0.29 | 7.36e-14 | 0.978 |
| LIHC | pam | 4 | 0.29 | 0.008 | 0.984 |
| LUSC | pam | 5 | 0.39 | 0.02044 | 0.969 |
| OV | pam | 3 | 0.31 | 0.00254 | 0.992 |
| SKCM | pam | 5 | 0.46 | 6.56e-8 | 0.988 |
| STAD | pam | 3 | 0.48 | 0.00961 | 0.99 |
| UCEC | pam | 2 | 0.6 | 0.00727 | 0.996 |

**Figure 2.**
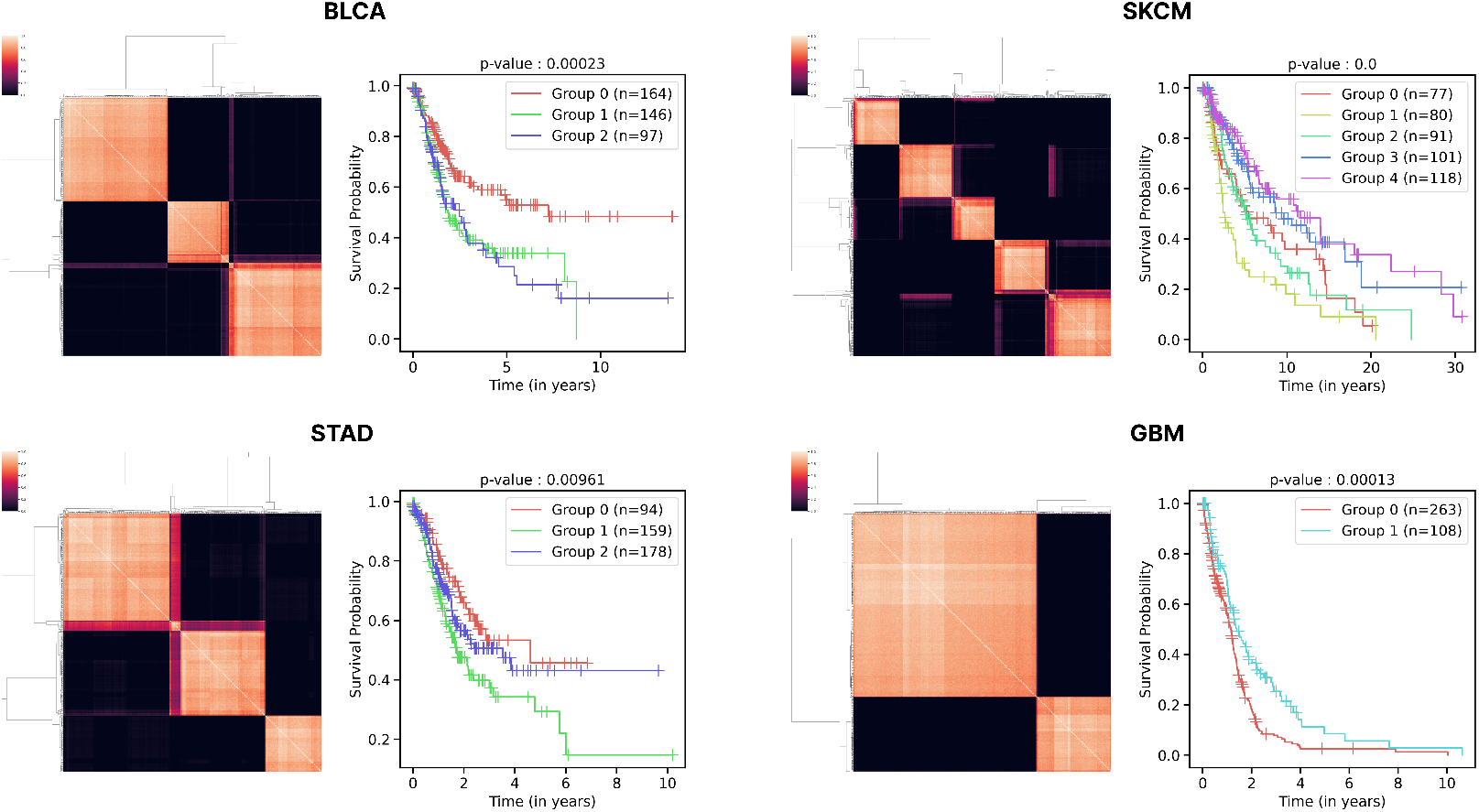
Unsupervised clustering for four cancer types: BLCA (Bladder Urothelial Carcinoma), STAD (Stomach Adenocarcinoma), SKCM (Skin Cutaneous Melanoma), and GBM (Glioblastoma Multiforme). On the left, the consensus map illustrates the clustering stability and agreement across multiple runs, visually representing the robustness of the clusters formed. On the right, Kaplan-Meier plots are displayed for each cancer type, offering insights into the survival probabilities associated with the clusters identified.

### Graph-learning based survival prediction model

We also adopted a supervised approach, which involved training a Cox regression model on the embeddings using patient survival data. This model was used for survival prediction and the model performance was evaluated using the c-index value. The c-index varied across cancer types, ranging from 0.5 to 0.8. For BRCA, CESC, COAD, KIRC, KIRP, LIHC, and LGG, we obtained a c-index above 0.7, indicating high hazard-predictive capability. For BLCA, LUAD, LUSC, HNSC, SKCM, STAD, and UCEC, the c-index ranged between 0.6 and 0.7, reflecting good performance in hazard prediction task(Table 2). These results demonstrates the model’s robust performance across various cancer types. We also divided patients into 4 groups based on Hazard scores obtained from Cox-model. These four groups showed significant survival differences for all 16 cancer types. Patients in lower hazard score percentiles exhibited significantly better survival rates than those in higher percentiles (Figure 3 and Figure S2).

**Table 2.** Supervised learning using graph-based embeddings: The c-index obtained for the Cox hazard prediction task on the encodings obtained using the DGM pipeline with NCG network is given.

| Cancer Type | C-Index |
| --- | --- |
| BLCA | 0.64 |
| BRCA | 0.72 |
| CESC | 0.78 |
| COAD | 0.74 |
| GBM | 0.56 |
| HNSC | 0.66 |
| KIRC | 0.73 |
| KIRP | 0.73 |
| LGG | 0.8 |
| LIHC | 0.71 |
| LUAD | 0.63 |
| LUSC | 0.62 |
| OV | 0.59 |
| SKCM | 0.61 |
| STAD | 0.64 |
| UCEC | 0.69 |

**Figure 3.**
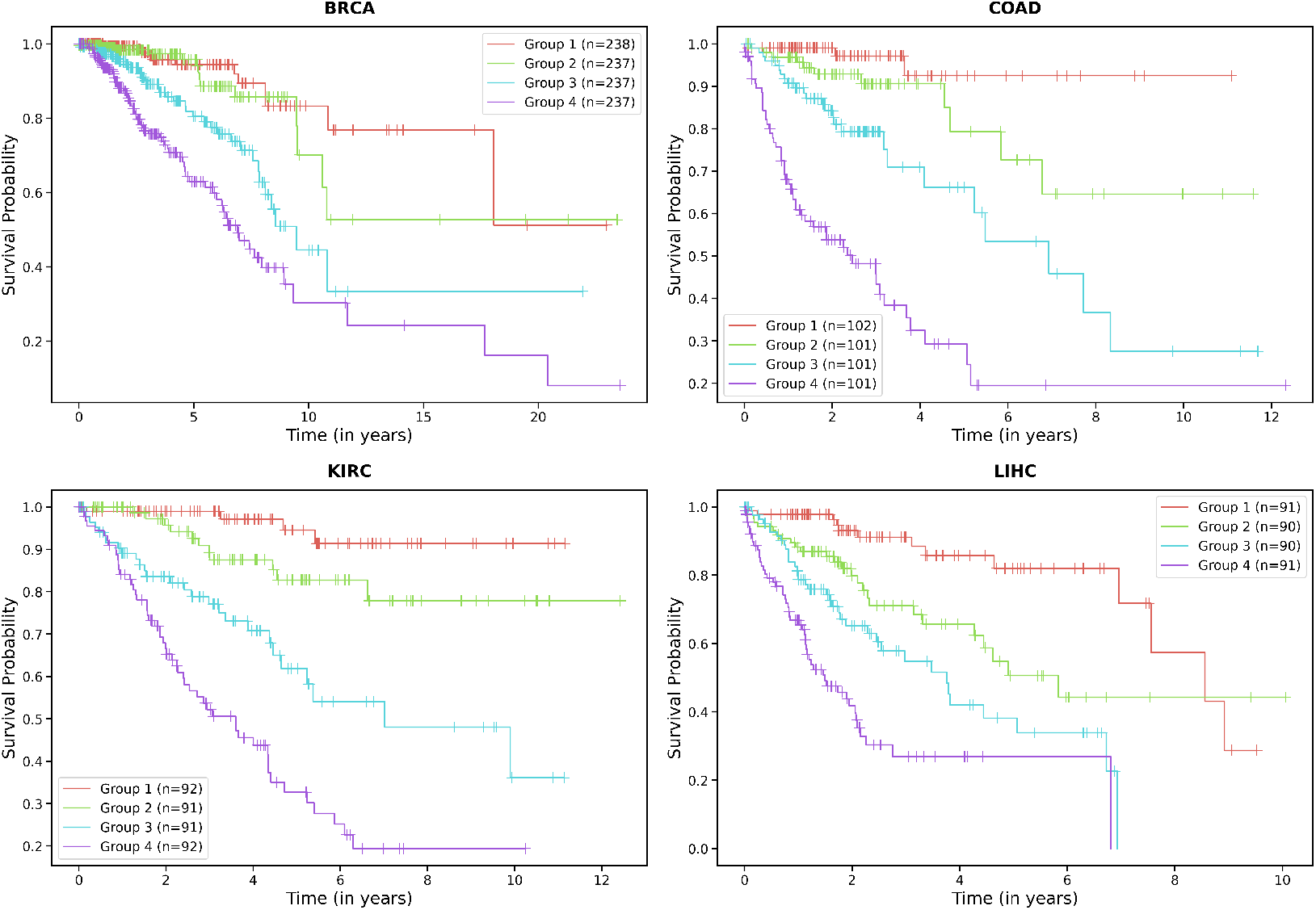
Kaplan-Meier (KM) plots for patients stratified into four groups according to the quartiles of hazard scores predicted by our Cox proportional hazards model with the cancer types BRCA (Breast Invasive Carcinoma), COAD (Colon Adenocarcinoma), KIRC (Kidney Renal Clear Cell Carcinoma), and LIHC (Liver Hepatocellular Carcinoma). Each group in the plot represents one of the quartiles.

### Robustness of encoders

To evaluate the impact of attention mechanisms on encoder performance, we repeated our experiments using a simplified, lightweight encoder based on the SAGEConv layer, termed DGMS. The unsupervised clustering and survival analysis revealed significant survival differences between patient groups for DGM compared to DGMS, as evident from the corresponding p-value (Table S2). Unsupervised clustering of DGMS encodings yielded significant results in 10 out of the 16 cancers studied. However, DGM clusters exhibited superior separation compared to those of DGMS, which is evident from the silhouette score and the log-rank p-value. In supervised analysis, the DGMS model encodings achieved a c-index over 0.7 for only 4 cancers and between 0.6 and 0.7 for 7 cancers. In contrast, the DGM model outperformed DGMS, achieving c-index values above 0.7 for 7 cancers (Figure S3).

In our experiments, both autoencoder models generated relevant encodings across the majority of the cancer types. Notably, the DGM model exhibited superior robustness and higher consistency in the clinical relevance of its encodings compared to its DGMS counterpart. Moreover, this graph attention-based model displayed a faster convergence rate during the training phase and achieved a lower minimum in the loss function, indicating a more efficient learning process. However, it is crucial to acknowledge the increased computational complexity introduced by the graph attention mechanism. This complexity not only necessitates more computational resources but also leads to longer training durations compared to the model without attention mechanisms.

### Network comparison

The additional networks were used to investigate model performance with different network sizes and node/edge compositions. We compared the results obtained using the NCG network with another reported-cancer-specific network, CancerReferenceNetwork (CRN)^21^. Due to the usage of different gene sets for the creation of the networks, there are significant disparities in the nodes (genes) and edges encompassed by each network (NCG and CRN) (Figure S4). The CRN yielded significant performance for 10 cancers (Table S3), while the unified network of CRN and NCG achieved notable results in 11 of the 16 cancer types (Table S4). However, their p-values were not as good as those of NCG-based encodings in most cases (Table S2). These networks also showed cancer-specific differences in performance. A notable case is observed in the case of KIRC, which was not accurately clustered using the CRN. This evidence suggests that different networks capture distinct, cancer-specific feature landscapes, emphasizing the importance of selecting the appropriate network to accurately interpret the complex biological information presented by omics data. It was observed that an increase in network size correlated with a reduction in clustering stability and a rise in computational complexity. This complexity was primarily attributed to the expansion of the attention matrix within the computational process despite the number of the model’s learnable parameters remaining unchanged. An evaluation of the simpler DGMS model, which lacks an attention mechanism, demonstrated satisfactory outcomes in 9 out of 16 cases with the NCG network, 6 out of 16 cases with the CRN, and 8 out of 16 cases with the Unified Network. Additionally, the Cox model performance on the encodings revealed that the NCG and Unified network-based encodings achieved higher c-index values compared to the CRN (Figure S5). These findings highlight the balance between network complexity, computational requirements, and clustering efficacy, with the NCG network emerging as the most favourable option across these dimensions in the context of cancer subtype identification.

### Performance comparison with network-based stratification method

Our framework focuses on graph-level task, distinguishing it from most deep learning approaches in this domain, which have primarily focused on multi-omics data integration^17^. Given the differences in data types, direct comparison with these methods may not be appropriate. To ensure a fair and rigorous evaluation, we initially benchmarked our method using the SAGEConv encoder (Table S2 and Figure S3), a widely recognized model in genomic research. Following this, we now compare our framework to the well-established PyNBS method^13^, which uses network propagation with somatic mutation data to encode PPI networks and generate patient representations. PyNBS applies non-negative matrix factorization (NMF) followed by hierarchical clustering to categorize patients into subtypes. PyNBS serves as a relevant benchmark due to its conceptual similarity to our approach: both frameworks propagate somatic mutation data across a network to perform cancer subtyping, making it a suitable reference for assessing our model’s performance. The PyNBS model was applied to the 16 cancers considered in our study using the NCG network to allow for direct comparison. Figure 4 compares the performance of PyNBS and DGM in unsupervised tasks, reporting p-values and cophenetic correlation coefficients. PyNBS yielded significant p-values for only 3 cancer types, whereas DGM achieved significance for 11. Figure 5 compares their performance in supervised tasks using the c-index, where DGM consistently outperformed PyNBS. These results demonstrate the superior encoding capability of our DGM model over PyNBS in both supervised and unsupervised downstream tasks.

**Figure 4.**
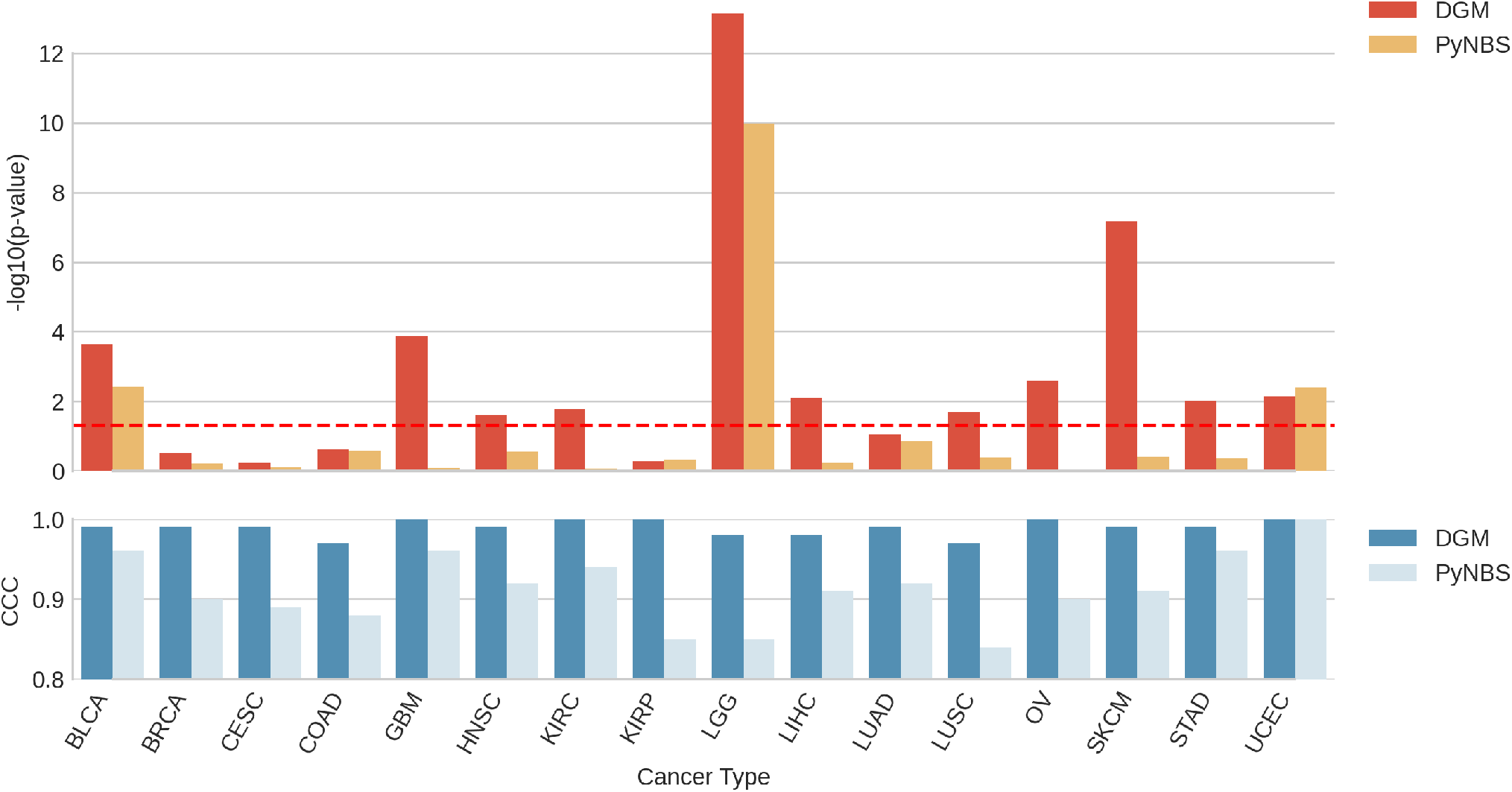
Comparative analysis with PyNBS in unsupervised task.

**Figure 5.**
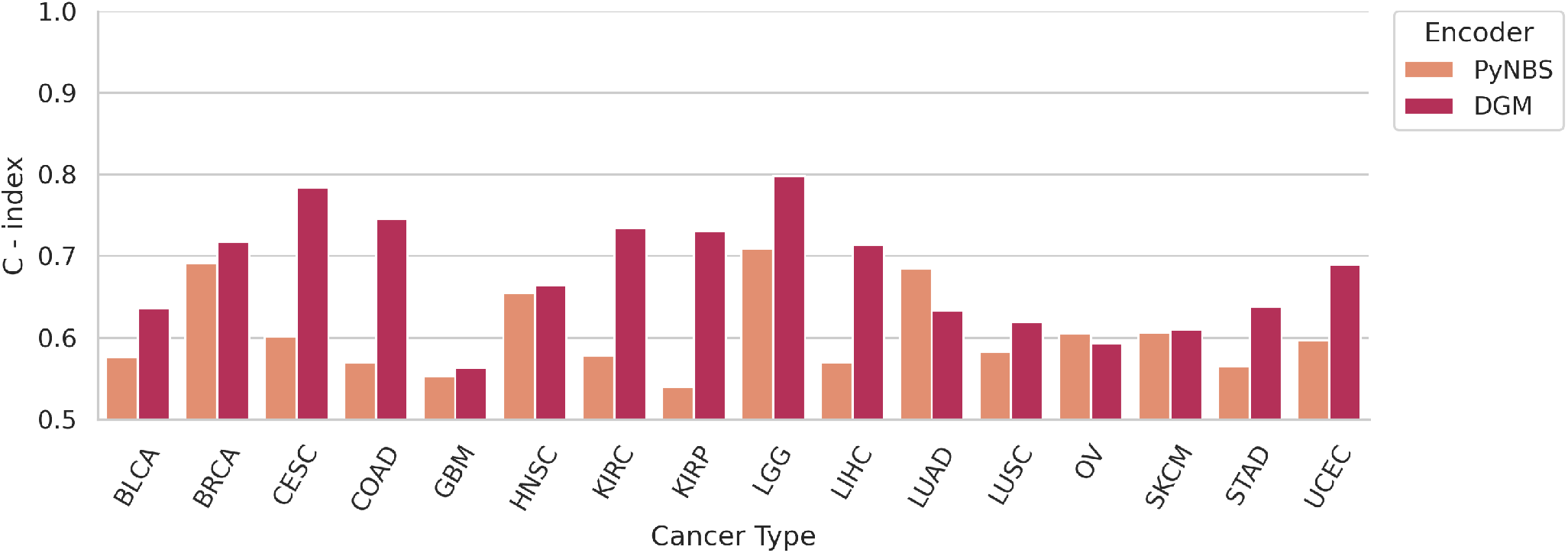
Comparison of performance of DGM and PyNBS in supervised task.

## Discussion

Somatic mutation profiling is becoming important for precision diagnostics and treatment. Heterogeneity in somatic mutation profile of specific cancers raises the challenge in accurately stratifying patients for cancer prognosis and treatment. Biological network provides an alternative view to molecular world. In this study, we have proposed a novel graph-based deep learning model called DGM for integrating patient mutation profile with prior network information for cancer subtype identification and prognosis across multiple cancer types.

The proposed DGM pipeline utilize the attention mechanisms in both the encoder and decoder, which set our method apart from regular graph autoencoding approaches The encoder’s attention focuses on crucial edges in the PPI network, while the decoder’s node-level attention enables effective decoding of node values. The utilisation of graph normalisation techniques in the pipeline helps to process the gene embeddings effectively during the encoding process. This also led to the faster convergence of the loss function during the training process. The use of focal loss addresses the sparsity of the dataset, ensuring accurate prediction of the presence and absence of somatic mutations. In the unsupervised analysis, DGM consistently identified significant clusters across 11 of the 16 cancers considered. The supervised analysis revealed a high correlation between the encodings and survival outcomes for all 16 cancer types. The integration of the NCG network yielded superior encodings compared to other networks used in the study, likely due to the presence of more relevant nodes and edges specific to cancers. Furthermore, the PAM clustering algorithm generated clusters with balanced patient distribution and clear separation of survival outcomes.

Our methodology extends the previously established NBS approach, which has been successfully applied in multiple studies using different prior networks and omics data types to obtain clinically relevant encodings^14^. The DGM pipeline improves upon the existing NBS framework and help to stratify various cancer types using a generic cancer-specific network and graph-learning paradigm. Importantly, our single-omics approach achieves c-index values that are comparable to or better than those reported in multi-omics studies for the same cancer types^22^. We also simplified the pipeline by employing a lightweight model DGMS (SAGEConv encoder), which achieved reasonable performance across a few cancer types. By focusing solely on somatic mutation data within the network of cancer-related genes, our models reduce computational complexity and offer a more streamlined solution.

The main limitations of our study include the inability of the model to reduce the encoding size, which will always remain as the number of nodes in the network, and the incompleteness of the PPI networks and gene sets used. A smaller, more relevant gene set, along with a more accurate interaction network, could potentially yield smaller, more meaningful encodings. Our experiments indicate that the network size does not directly correlate with clinical outcomes. Instead, gene sets, and interaction lists (edges) used play a significant role. Additionally, the current model has only been tested and tuned on somatic mutation profiles, and its ability to encode other omics data has not been explored. In conclusion, the proposed models show promising results in capturing clinically relevant information from somatic mutation data, even in the presence of high sparsity levels. These findings highlight the potential of our approach for cancer subtype identification and prognosis in resource-limited clinical diagnostic settings, where multi-omics data may not be readily available.

## Methods

### Omics Data

Somatic mutation data for 16 distinct cancer types, along with the corresponding clinical information, were systematically retrieved from the Genomic Data Commons (GDC)^23^ data portal using the TCGABiolinks (2.25.0) R package^24^. Only cancers with a cohort size exceeding 250 patient samples were considered. The somatic mutation data was then processed into patient-by-gene matrices, with “0” indicating the absence of a mutation and “1” indicating its presence. It is important to note that these matrices are over 95% sparse.

### Network Assembly

The primary network, was constructued using cancer genes (altered genes driving cancer) and healthy drivers (altered genes driving noncancer clones) from the network of cancer-related genes database^20^. We then isolated the edges that interconnect the gene set obtained from the database using PCNet, a parsimonious network compiled by Huang et al (2018)^21^. This process resulted in the creation of the NCG Network. Additionally, we utilized two other networks to examine the impact of network choice on performance. The first network, referred as the CancerReference Network (CRN), included genes used in the PyNBS study^13^, with edges extracted from PCNet. The second network referred to as the “Unified Network,” was constructed by combining the edge lists from both the CRN and NCG Networks.

### Model Architecture Encoder

This study proposes two encoder designs, each featuring the same backbone architecture but employing different graph layers (Figure 1). The first encoder, DGM, integrates graph attention layers, specifically employing a graph layer called TransformerConv^25^. In contrast, the second model, DGMS, utilizes the SAGEConv graph layer^26^ with a mean aggregator. The incorporation of graph attention layers enables the model to prioritize crucial edges by computing attention weights, thereby enhancing the encoding process. Additionally, GraphNorm^27^, a normalization strategy that leverages graph structure information, is applied to optimize the normalization procedure. This strategy presents significant advantages over traditional normalization techniques, particularly in handling heterogeneous data types and complex network structures. The combination of these methods enhance the overall encoding process, leading to more robust and accurate representations of graph-structured data. For subsequent downstream analysis, the mean pooled output from the encoder layer was used to generate a feature list representing gene-wise information. The message-passing in the TransformerConv and the SAGEConv layers is governed by the following equations:

TransformerConv:

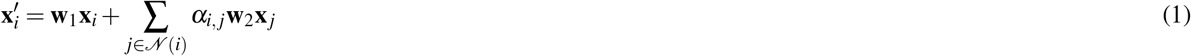

where the attention coefficients α_i, j_ are computed via multi-head dot product attention:

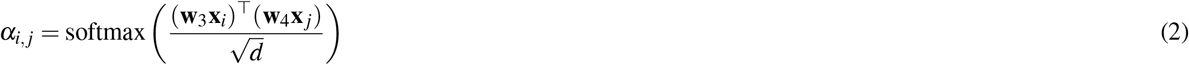

α_i, j_ = softmax SageConv:

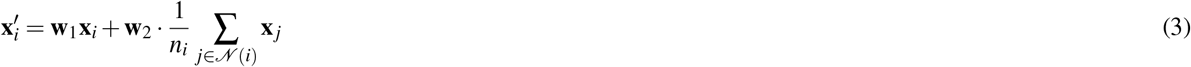

Here, *x_i_* and *x_j_* denote the feature vectors of node *i* and its neighbour *j*, respectively, while *x_i_^′^* represents the updated feature vector of node *i* after applying the convolution. The learnable weight matrices *w*_1_ and *w*_2_ transform the features of the central node and its neighbours, while *w*_3_ and *w*_4_ are used to compute the attention coefficients *α_i,j_*, which determine the importance of neighbour *j* when updating the features of node *i* in the TransformerConv layer. The dimensionality of the feature vectors is represented by *d*, and the softmax function is used to normalize the attention coefficients. *𝒩(i)* represents the set of neighbours of node *i*, and *n_i_* is the number of neighbours used for normalization in SageConv.

### Decoder

We propose a node decoder architecture with an integrated multi-head attention mechanism to overcome the limitations of traditional graph autoencoders that use edge decoders^28^. Traditional edge decoders generate an adjacency matrix by performing an inner product on the encodings, which is not suitable for our specific application. In our analysis, the graph structure remains consistent across all patient samples, making the adjacency matrix invariant and limiting the effectiveness of edge decoders in capturing discriminative representations. In contrast, our attention-augmented node decoder is designed to effectively capture node features while accommodating the static nature of the edge list across different patient samples.

### Loss Function

We used Focal Loss^29^ to address the challenge of handling highly sparse node feature data. Focal loss is a modification of cross-entropy loss that allows for the modulation of importance given to 0s and 1s within the dataset through the adjustment of the hyperparameter gamma. The formulation of loss function is detailed in Equation 4. Traditional loss functions such as cross-entropy and mean squared error (MSE) prove to be ineffective for our datasets as they predominantly account for the presence of mutations (1s) without adequately addressing the absence of mutation (0s). By employing Focal Loss, we enhance the model’s ability to prioritize and accurately classify both the 0s and 1s, thereby improving the model’s performance on our sparse dataset.

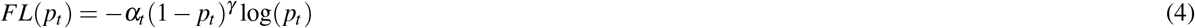

In the given equation, (*p_t_*) signifies the probability predicted for the actual class. The parameter (*α_t_*) acts as a balancing factor between positive and negative instances, often assigned the value of the inverse class frequency. Additionally,(*γ*) serves as a focusing parameter.

### Model Hyperparameter Tuning and Training

In this study, 80% of the data was used for training and the rest 20% for validation. The optimal output channel size was identified as 10 through grid search analysis, which revealed that increasing the number of channels beyond this point did not significantly reduce the loss, and a smaller channel size would lead to oversquashing^30^. We set the learning rate at 1e-4 after testing to optimize model performance without compromising efficiency. The Gamma parameter in the focal loss was set to 2. The Adam optimizer was chosen for its effectiveness in optimizing the loss function and training was expedited using the PyTorch Lightning and PyToch Geometric frameworks. The number of parameters in both the encoder and decoder depends solely on the number of output channels considered in the study and remains constant across all cancers and networks considered. Experiments were conducted on NVIDIA GTX 2080Ti GPUs over 200 epochs. To optimize resource use, we employed a batch size of 2, accumulating gradients over 4 batches before backpropagation.

### Unsupervised Analysis of encodings

In our study, we used a consensus clustering framework to analyze the encodings from the graph autoencoder, with the goal of achieving robust and reproducible clustering results. This approach is implemented through two distinct clustering algorithms: the PAM algorithm^31^, which is effective in handling outliers by selecting actual data points as cluster centers, and the K-means algorithm^32^, which offers computational efficiency for large datasets despite being sensitive to initial conditions. The adoption of consensus clustering^33^ serves to mitigate the variability inherent in clustering outcomes, ensuring that the clusters identified are stable and consistent across different iterations of the experiment.

The evaluation of cluster effectiveness and significance in this study was based on three critical metrics. Firstly, the cophenetic correlation coefficient was used to asses the integrity of the consensus clustering. This coefficient measures the correlation between the original distances among data points and the distances represented in the clustering dendrogram, providing insight into how well the clustering solution reflects the original dataset’s structure. A higher cophenetic correlation coefficient indicates that the clustering arrangement more accurately represents the inherent relationships among data points, thus validating the effectiveness of consensus clustering. Secondly, the Silhouette score was calculated to evaluate how well the patients fit into the predicted clusters^34^. Lastly, the log-rank test^35^ was used as the primary method to assess differences in survival probabilities across clusters. We applied a p-value threshold of < 0.05 to identify significant clusters. These survival analyses were performed using the ‘lifelines’ package in Python^36^.

### Supervised Analysis of encodings

The Cox proportional hazards model was trained using the encodings for survival prediction^37^. We employed a split-sample methodology(70:30) and performed a 5-fold cross-validation 100 times. We reported the mean C-Index, which is a reliable indicator of the model’s performance. We also calculated the hazard score for each patient and categorized them into four groups based on the quantile hazard ratios. Survival differences between these groups were analyzed using Kaplan-Meier plot.

## Acknowledgements

This work was supported by iHUB-Data, International Institute of Information Technology, Hyderabad, India.

## Author contributions statement

Conceptualization: P. K. Vinod; Methodology: Aswin Jose; Formal analysis and investigation: Aswin Jose, Akansha Srivastava; Writing - original draft preparation: Aswin Jose, Akansha Srivastava; Writing - review and editing:Aswin Jose, Akansha Srivastava, P. K. Vinod; Funding acquisition: P. K. Vinod; Supervision: P. K. Vinod

## References

1. Bray, F. et al. Global cancer statistics 2022: Globocan estimates of incidence and mortality worldwide for 36 cancers in 185 countries. CA: a cancer journal for clinicians 74, 229–263 (2024).

2. Carbone, A. Cancer classification at the crossroads. Cancers 12, 980 (2020).

3. Vargo-Gogola, T. & Rosen, J. M. Modelling breast cancer: one size does not fit all. Nat. Rev. Cancer 7, 659–672 (2007).

4. Kulavi, S., Ghosh, C., Saha, M. & Chatterjee, S. One size does not fit all: An overview of personalized treatment in cancer. J. Pharm. Res. Int. 87–103 (2021).

5. Cooper, G. M. The Cell: A Molecular Approach (Sinauer Associates, Sunderland (MA), 2000), 2nd edn.

6. Reuter, J. A., Spacek, D. V. & Snyder, M. P. High-throughput sequencing technologies. Mol. cell 58, 586–597 (2015).

7. Carter, H. et al. Cancer-specific high-throughput annotation of somatic mutations: computational prediction of driver missense mutations. Cancer research 69, 6660–6667 (2009).

8. Barabasi, A.-L. & Oltvai, Z. N. Network biology: understanding the cell’s functional organization. Nat. reviews genetics 5, 101–113 (2004).

9. Redhu, N. & Thakur, Z. Network biology and applications. Bioinformatics 381–407 (2022).

10. Zhang, P. & Itan, Y. Biological network approaches and applications in rare disease studies. Genes 10, 797 (2019).

11. Hofree, M., Shen, J. P., Carter, H., Gross, A. & Ideker, T. Network-based stratification of tumor mutations. Nat. methods 10, 1108–1115 (2013).

12. Zhong, X., Yang, H., Zhao, S., Shyr, Y. & Li, B. Network-based stratification analysis of 13 major cancer types using mutations in panels of cancer genes. BMC genomics 16, 1–8 (2015).

13. Huang, J. K., Jia, T., Carlin, D. E. & Ideker, T. pynbs: a python implementation for network-based stratification of tumor mutations. Bioinformatics 34, 2859–2861 (2018).

14. Di Nanni, N., Bersanelli, M., Milanesi, L. & Mosca, E. Network diffusion promotes the integrative analysis of multiple omics. Front. genetics 11, 488641 (2020).

15. Liu, C., Han, Z., Zhang, Z.-K., Nussinov, R. & Cheng, F. A network-based deep learning methodology for stratification of tumor mutations. Bioinformatics 37, 82–88 (2021).

16. Xia, F. et al. Graph learning: A survey. IEEE Transactions on Artif. Intell. 2, 109–127 (2021).

17. Wekesa, J. S. & Kimwele, M. A review of multi-omics data integration through deep learning approaches for disease diagnosis, prognosis, and treatment. Front. Genet. 14, 1199087 (2023).

18. Liang, C., Shang, M. & Luo, J. Cancer subtype identification by consensus guided graph autoencoders. Bioinformatics 37, 4779–4786 (2021).

19. Baul, S., Ahmed, K. T., Filipek, J. & Zhang, W. omicsgat: Graph attention network for cancer subtype analyses. Int. J. Mol. Sci. 23, 10220 (2022).

20. Repana, D. et al. The network of cancer genes (ncg): a comprehensive catalogue of known and candidate cancer genes from cancer sequencing screens. Genome biology 20, 1–12 (2019).

21. Huang, J. K. et al. Systematic evaluation of molecular networks for discovery of disease genes. Cell systems 6, 484–495 (2018).

22. Cheerla, A. & Gevaert, O. Deep learning with multimodal representation for pancancer prognosis prediction. Bioinformatics 35, i446–i454 (2019).

23. Grossman, R. L. et al. Toward a shared vision for cancer genomic data. New Engl. J. Medicine 375, 1109–1112 (2016).

24. Huber, W. et al. Orchestrating high-throughput genomic analysis with bioconductor. Nat. methods 12, 115–121 (2015).

25. Shi, Y. et al. Masked label prediction: Unified message passing model for semi-supervised classification. In Proceedings of the Thirtieth International Joint Conference on Artificial Intelligence, 1548–1554, DOI: 10.24963/ijcai.2021/214 (ijcai.org, 2021).

26. Hamilton, W., Ying, Z. & Leskovec, J. Inductive representation learning on large graphs. Adv. neural information processing systems 30 (2017).

27. Cai, T. et al. Graphnorm: A principled approach to accelerating graph neural network training. In International Conference on Machine Learning, 1204–1215 (PMLR, 2021).

28. Kipf, T. N. & Welling, M. Variational graph auto-encoders. arXiv preprint 1611.07308 (2016).

29. Lin, T.-Y., Goyal, P., Girshick, R., He, K. & Dollár, P. Focal loss for dense object detection. In Proceedings of the IEEE international conference on computer vision, 2980–2988 (2017).

30. Alon, U. & Yahav, E. On the bottleneck of graph neural networks and its practical implications. In International Conference on Learning Representations (2021).

31. Park, H.-S. & Jun, C.-H. A simple and fast algorithm for k-medoids clustering. Expert. systems with applications 36, 3336–3341 (2009).

32. Jin, X. & Han, J. K-means clustering. Encycl. machine learning 563–564 (2011).

33. Strehl, A. & Ghosh, J. Cluster ensembles—a knowledge reuse framework for combining multiple partitions. J. machine learning research 3, 583–617 (2002).

34. Rousseeuw, P. J. Silhouettes: a graphical aid to the interpretation and validation of cluster analysis. J. computational applied mathematics 20, 53–65 (1987).

35. Kaplan, E. L. & Meier, P. Nonparametric estimation from incomplete observations. J. Am. Stat. Assoc. 53, 457–481, DOI: 10.1080/01621459.1958.10501452 (1958).

36. Davidson-Pilon, C. lifelines: survival analysis in python. J. Open Source Softw. 4, 1317, DOI: 10.21105/joss.01317 (2019).

37. Cox, D. R. Regression models and life-tables. J. Royal Stat. Soc. Ser. B (Methodological) 34, 187–202 (1972).

